# Maize mutant hybrids with improved drought tolerance and increased yield in a field experimental setting

**DOI:** 10.1101/2025.07.10.664191

**Authors:** Federico Belén, Paloma Garnero Patat, Camila Jaime, Santiago Walker, Ignacio Dellaferrera, José Maiztegui, Germán Dunger, Marcela Dotto

## Abstract

Previous studies determined that maize mutants in miR394-regulated genes, ZmLCR1 and ZmLCR2, are more tolerant than wild-type seedlings to prolonged periods of drought. In order to evaluate the effect of these mutations in a genetic background more similar to that of maize commercialization, in this work we evaluate growth of double mutant hybrid plants in W22/B73 genetic background and also evaluated plant fitness, flowering and yield in experimental plots under two watering regimes, and compared the nutritional content of wild-type and mutant hybrids. Our results show that mutant hybrid seedlings exhibit improved physiology under normal watering conditions as well as in drought conditions, exhibiting an increase in epicuticular wax content, unaltered membrane damage in drought and lower ROS production, supporting higher survival after severe drought for double mutant hybrid seedlings. We also established that the hybrid mutants grown in typical agricultural conditions do not show differences in flowering time or in physiological and nutritional aspects, but they present a higher yield in comparison to wild-type W22/B73 hybrids, as determined by higher ear weight and number of kernels per ear in mutant hybrids, when grown in field rainfed conditions.

**Highlights:**

- Double mutants in miR394-regulated genes, ZmLCR1 and ZmLCR2, show enhanced drought tolerance in hybrid maize seedlings, with improved physiological traits under both normal and stress conditions.
- Mutant hybrids exhibit increased epicuticular wax accumulation and reduced ROS production, supporting greater survival during prolonged drought stress.
- Field-grown mutant hybrids in a W22/B73 background maintain normal flowering time and nutritional composition, indicating no agronomic penalties from the mutations.
- Yield is significantly higher in mutant hybrids compared to wild-type controls, as shown by increased ear weight and kernel number under rainfed field conditions.
- These findings highlight ZmLCR1 and ZmLCR2 as valuable targets for breeding drought-tolerant, high-yielding maize cultivars suited to production environments.

## 1. Introduction

Abiotic environmental factors such as extreme temperatures, salinity and drought significantly impact plant growth and development. To cope with these adverse conditions, plants must rapidly activate coordinated responses, modulating the expression and activity of various molecular networks (Chirivì and Betti, 2023; Zhang et al., 2023).

Drought, defined as a prolonged water deficit that impairs plant growth and survival, is a major challenge in agriculture. It alters key physiological traits and can ultimately reduce crop yields (Martignago et al., 2020). Additionally, water scarcity affects soil nutrient availability and transport, as water serves as the primary medium for nutrient movement (Chaudhry and Sidhu, 2022). The economic impact of drought is exacerbated by climate change and decreasing rainfall patterns (Kim et al., 2019).

In maize cultivation, water requirements vary across developmental stages. While early vegetative growth demands minimal water, the need peaks during the reproductive phase before declining in later stages. Drought stress during vegetative growth slows maize development, extending the vegetative phase and shortening the reproductive stage (Bheemanahalli et al., 2022; Pandian et al., 2020). Notably, maize is highly sensitive to water deficits and high temperatures during flowering due to its cross-fertilizing nature. Both pollen and stigmas are directly exposed to environmental conditions, making them vulnerable to drought-induced damage. Key symptoms include delayed inflorescence development, floral asynchrony, reduced pollen fertility, and impaired grain filling, all of which significantly impact reproductive success and yield (Bheemanahalli et al., 2022; Chirivì and Betti, 2023; Salehi-Lisar and Bakhshayeshan-Agdam, 2020).

Water availability is crucial for metabolic processes, including photosynthesis, since drought-induced stomatal closure limits COL uptake and chlorophyll degradation reduce photosynthetic efficiency (Lawlor and Tezara, 2009; Mundim and Pringle, 2018). Consequently, physiological parameters such as photosynthetic rate, transpiration, stomatal conductance, and chlorophyll content are commonly used to assess plant water status. In addition, stress conditions induce reactive oxygen species (ROS) production, leading to activation of ROS-scavenging mechanisms and downstream signaling to cope with the challenging environment (Samanta et al., 2024; Wang et al., 2024). Hence, some drought-tolerant species enhance water-use efficiency by minimizing water loss, a key adaptation for survival under water-limited conditions (Bramley et al., 2013), while genes involved in ROS homeostasis have been proposed as key candidates to develop crops with enhanced tolerance to abiotic stress in plants (Nadarajah, 2020). Conversely, excess moisture in maize production, caused by flooding, waterlogging or poor soil drainage, can also be an important constraint in some tropical and rainy growing areas. When plants lack a proper ventilation system for gaseous exchange between aboveground plant parts and inundated roots, they suffer with extreme oxygen stress, hypoxia followed by anoxia, whenever it faces prolonged excess soil moisture situation (Zaidi et al., 2004; Thapa et al., 2025).

In maize breeding, classical genetic improvement has been instrumental in developing inbred lines for hybrid production, a cornerstone of modern maize cultivation. The exploitation of heterosis, where hybrid offspring outperform parental lines in growth and productivity, has led to significant yield gains (Benavente and Giménez, 2021; Chen, 2010; Comings and MacMurray, 2000). Hybrid maize exhibits superior grain production efficiency and enhanced tolerance to biotic and abiotic stresses, making it widely adopted in agriculture (Li et al., 2021; Xiao et al., 2021). Beyond classical breeding, biotechnology has enabled the development of genetically modified (GM) crops, initially through transgenesis and, more recently, via genome editing technologies (Dong et al., 2019; Klümper and Qaim, 2014; Simmons et al., 2021). GM crops have demonstrated success in mitigating environmental stress impacts, improving productivity, and benefiting farmers (Klümper and Qaim, 2014; Que et al., 2014).

Recent studies in maize have identified the miR394 pathway as key regulator of drought tolerance. This microRNA targets the ZmLCR1 and ZmLCR2 genes, orthologs of *LEAF CURLING RESPONSIVENESS (LCR)* in Arabidopsis. Mutant lines lacking ZmLCR1 and ZmLCR2 exhibit enhanced drought tolerance, with a higher survival rate (84–92%) compared to wild-type (WT) plants (35%) after re-watering. These mutants maintain higher photosynthetic activity, improved intrinsic water-use efficiency, increased secondary root development, and elevated epicuticular wax production under drought conditions (Miskevish et al., 2025).

In the present study, we investigate the effects of these mutations in a hybrid W22/B73 genetic background. We assess seedling growth under drought stress and, in field conditions, we analyze plant fitness, flowering, and yield under different watering regimes, evaluating also the nutritional content of mutant and wild-type hybrids. This approach allows us to determine the potential benefits of these genetic modifications in a hybrid genetic background more representative of commercial maize production.

## 2. Materials and methods

### 2.1. Plant materials

Maize *zmlcr1* and *zmlcr2* single mutants, and *zmlcr1 zmlcr2* double mutants in W22 inbred background were obtained as previously described (Miskevish et al., 2025). To make hybrids, the *zmlcr1* and *zmlcr2* single mutations were introgressed into B73 background by back-crossing five times into B73 wild-type plants followed by self-pollination and identification of homozygous and heterozygous mutants for each gene (*zmlcr1/zmlcr1* and *zmlcr2/zmlcr2*; *zmlcr1/+* and *zmlcr2/+*). Next, *zmlcr1/zmlcr1* mutants were crossed to *zmlcr2/+* plants, followed by the identification of *zmlcr1/+ zmlcr2/+* individuals, which were next crossed to *zmlcr1/zmlcr1* mutants. The progeny was genotyped to identify *zmlcr1/zmlcr1 zmlcr2/+* individuals for self-pollination and identification of *zmlcr1/zmlcr1 zmlcr2/zmlcr2* double mutants in B73 background, which were finally bulked. To obtain hybrids, *zmlcr1/zmlcr1 zmlcr2/zmlcr2* double mutants in B73 were crossed to *zmlcr1/zmlcr1 zmlcr2/zmlcr2* double mutants in W22 background and plants from the F1 generation were used for all experiments, along with wild-type hybrids obtained by crossing W22 and B73 wild-type plants. For simplicity, wild-type hybrid plants will be referred to as W22/B73 and homozygous double mutant hybrid plants as *zmlcr1 zmlcr2^W22/B73^*.

### 2.2. Growing conditions

For experiments in growth chambers, maize seeds were treated with Carbendazim 0.4% v/v and sown in soil in 1 l pots, using a mixture of 75% soil and 25% Growmix Multipro substrate (Terrafertil) and transferred to a growth chamber with controlled growing conditions, in a long-day light regime (16 h light/8 h dark) and temperature between 24 and 28 °C.

When grown in experimental plots, seeds were treated with Carbendazim 0.4% v/v and sown in field plots at the Donnet Campus of the Faculty of Agricultural Sciences, from National University of Littoral (FCA-UNL, -31.44217386570492, - 60.94119040708018), in a plot of 300 m2, with rows separated by 52 cm and plant distance of 25 cm to achieve a density of 76.900 plant/ha. Plants were fertilized using granulated urea (150 kg/ha) and Surcostart (50 kg/ha) when plants were in V3-V4 stages; 350 ml/ha of cypermethrin (0.14 w/v) were used for insect control, applied as needed. Wild-type and mutant hybrids were sown in groups of 30 per genotype and treatment, determining 6 different sub-parcels surrounded by commercial hybrids (P2089 VYHR), planted to eliminate edge effect and to allow for extra pollen available for open pollination.

### 2.3. Differential irrigation treatments

For plants in growth chambers, control conditions consisted of irrigation to maximum field capacity every two days throughout the experiment. Maize plants gown under drought conditions received normal irrigation up to V3 stage and at this point irrigation was suspended for 30 days. Watering was resumed for 7 days, and plants that resumed growth and greenness were recorded each day to evaluate plant survival. We used a binary scoring system for survival (alive or dead) based on the presence of green tissue and overall plant vigor, with plants showing no green tissue (wilting) classified as dead.

For maize plants grown in experimental plots, two sections were established: one half was grown under rainfed conditions, receiving only rainwater, while the other half was grown with supplemented irrigation, receiving rainwater plus additional irrigation applied manually between rainfalls.

### 2.4. Determination of epicuticular wax content

Epicuticular wax extraction was adapted from Bewick et al. (1993). For each group of plants (genotypes W22/B73 or *zmlcr1/zmlcr2^W22/B73^*and control or drought treatments; n = 5 for each genotype/treatment combination), 15 cm measured from the distal part of the leaves were collected from each plant. The leaves were immersed (excluding 1 cm in the cut region) for 20 s in 50 ml of chloroform, contained in pre-weighted tubes (initial weight), and the total area of the submerged leaf was calculated using ImageJ software. The chloroform solution containing the wax was kept in a safety hood until the chloroform was completely evaporated and the tubes were weighed (final weight). The wax content was determined by weight difference (final weight - initial weight) and the wax content per unit area (mg/cm^2^) was calculated.

### 2.5. Measurement of physiological parameters

Different variables associated with plant physiological status were measured: greenness index using the SPAD 502 chlorophyll meter (Minolta), transpiration (E), stomatal conductivity (gs) and photosynthetic rate (Pn) using the CIRAS-2 meter (PP biosystems) and water use efficiency (EUA = Pn/gs) was calculated for at least 5 plants per genotype and treatment.

### 2.6. Detection of reactive oxygen species

The detection of hydrogen peroxide (H_2_O_2_) accumulation was carried out using 3,3’-diaminobenzidine (DAB; Sigma-Aldrich), following the protocol established by Jaime et al. (Jaime et al., 2024). Wild-type and mutant hybrid plants (W22/B73 and *zmlcr1/zmlcr2*^W22/B73^) were grown under drought and standard irrigation conditions as described above, and utilized for the analysis. For each genotype, three individual plants were randomly chosen and leaf samples were collected from each plant for evaluation. Leaf fragments were immersed in a 1 mg/ml DAB solution (Sigma Aldrich) in plastic tubes and incubated in the dark for 18 hours. Subsequently, the samples were decolorized by soaking in 96% ethanol (v/v) and maintained at 37 °C. From each leaf, five fragments were mounted for observation under a fluorescence microscope (Leica M205 FAC) at 40X magnification. The resulting images were analyzed using ImageJ Fiji software.

For each plant, five images were randomly selected for quantitative analysis. Stained areas were identified by applying the normalized threshold function and further processed using the "H DAB" vector from the Colour Deconvolution plugin in ImageJ Fiji. The specific RGB vector parameters used were: R: 0.2681, G: 0.5703, B: 0.7764.

### 2.7. Yield determination

Ears of hybrid maize, both wild-type W22/B73 and double mutants *zmlcr1/zmlcr2^W22/B73^*grown in experimental plots, divided in 6 sub-parcels as detailed above, were harvested 50 days after pollination and dried at 27 °C for one week. To determine the yield, the following parameters were analyzed: weight per ear (g), weight of 1000 grains (g) and number of grains per ear (n ≥ 10, by genotype and treatment).

### 2.8. Nutritional analysis

Wild-type W22/B73 and double mutant *zmlcr1/zmlcr2^W22/B73^*hybrids were harvested from the field after senescence, and the vegetative aerial parts of adult plants were used to evaluate nutritional content. Dry matter (DM) was determined by oven-drying at 60 °C until a constant weight was achieved. Total nitrogen content and corresponding crude protein (CP) content were determined using the Kjeldahl method, following procedures of the Association of Official Analytical Chemists (AOAC, 1990. Official Methods of Analysis of AOAC International). Neutral detergent fiber (NDF), acid detergent fiber (ADF), and acid detergent lignin (ADL) were determined according to the method described by Van Soest et al. (Van Soest et al., 1991) and ether extract (EE) was determined using the methodology described by Thiex et al. (Thiex et al., 2003).

### 2.9. Statistical analysis

For statistical analysis of the results, two- or three-way ANOVA tests were used after verification of normality, according to the Shapiro-Wilk test and equality of variances, according to the Levene test (Levene, 1960; Shapiro and Wilk, 1965). For survival curves, the Kaplan–Meier log-rank analysis was used, followed by Holm-Sidak method for pairwise comparisons, and for nutritional composition Student’s t-tests was used. Statistical evaluations were performed using R Studio and Microsoft Excel was used for graphical representation of the data.

## 3. Results

### 3.1. Mutant hybrid seedlings show improved physiological status than wild-type hybrids

To verify that mutants in W22/B73 hybrid background exhibit a drought response similar to that observed in mutants in W22 inbred background (Miskevish et al., 2025), we started our analysis by conducting a drought experiment for hybrid seedlings in growth chambers. Results indicate that W22/B73 and *zmlcr1/zmlcr2*^W22/B73^ seedlings show no statistical differences at the beginning of the drought treatment (D1) for any of the analyzed traits associated with plant physiology (Fig. 1), indicating that at this point of plant development, right after seedlings received normal irrigation, physiology for both wild-type and mutant hybrids is similar for the variables greenness index, leaf transpiration (E), stomatal conductivity (gs), net photosynthesis (Pn) and intrinsic water use efficiency (iWUE) (Fig. 1B-F, D1). After 15 days of drought treatment (D15), plants show typical signs of water withdrawal, with rolled-up leaf blades and loss of turgidity, but mutant hybrids exhibit a slightly better overall status (Fig. 1A, representative individuals). At this point mutant hybrids show improved values for Pn and iWUE in comparison to wild-type hybrids subject to drought treatment (Fig. 1E-F, D15, Drought). Interestingly, *zmlcr1/zmlcr2*^W22/B73^ seedlings also showed improved values compared to wild-type seedlings at D15 for E, gs, Pn and iWUE, also in control conditions, exhibiting lower E, higher gs, Pn and iWUE in comparison to wild-type hybrids (Fig. 1C-F, D15, Control), suggesting that the mutations in hybrid backgrounds confer benefits to hybrid plants also under normal watering conditions. Surprisingly, *zmlcr1/zmlcr2*^W22/B73^ seedlings showed a lower greenness index in control conditions, compared to wild-type hybrids (Fig. 1B, D15, Control), which could be related to lower chlorophyl content for these plants.

**Figure 1:**
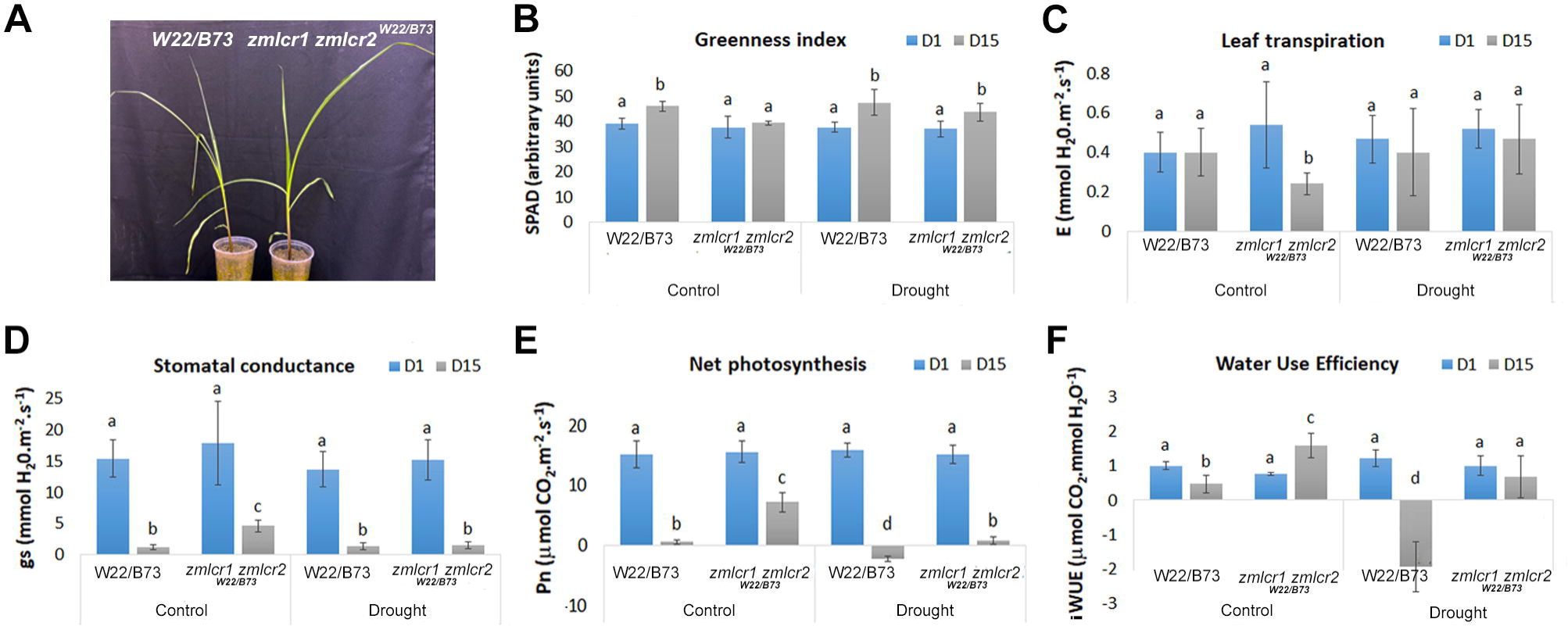
Physiological parameters of W22/B73 and *zmlcr1 zmlcr2^W22/B73^* hybrid seedlings. Measurements are shown for seedlings at the beginning of the drought treatment (D1) and after 15 days of drought (D15). A) Representative individuals at D15. Analysis of B) Greenness index, C) Leaf transpiration (E), D) stomatal conductance (gs), E) Net photosynthesis (Pn) and F) intrinsic Water Use Efficiency (iWUE) are shown, under normal irrigation (Control) and under drought conditions (Drought). Mean ± SD is represented; different letters indicate significant differences according to ANOVA (p < 0.05), using genotype, treatment and days as factors, and Tukey’s post-hoc tests for multiple comparisons; n ≥ 5.

### 3.2. Mutant hybrid seedlings are more tolerant to drought treatment

In order to evaluate factors that could be related to the improved physiology observed in hybrid seedlings, we analyzed epicuticular wax and ion leakage of plants after 15 days of drought treatment (D15). We determined that mutant hybrid seedlings have a higher content of epicuticular wax in drought conditions, whereas for wild-type hybrid seedlings the amount of wax remains unchanged (Fig. 2A). Moreover, wild-type hybrids exhibit the expected increased ion leakage in drought conditions, but for mutant hybrids no statistically significant difference was observed (Fig. 2B). These results indicate that these factors could be related to the improvement in physiological parameters observed for mutant hybrids in drought conditions.

**Figure 2:**
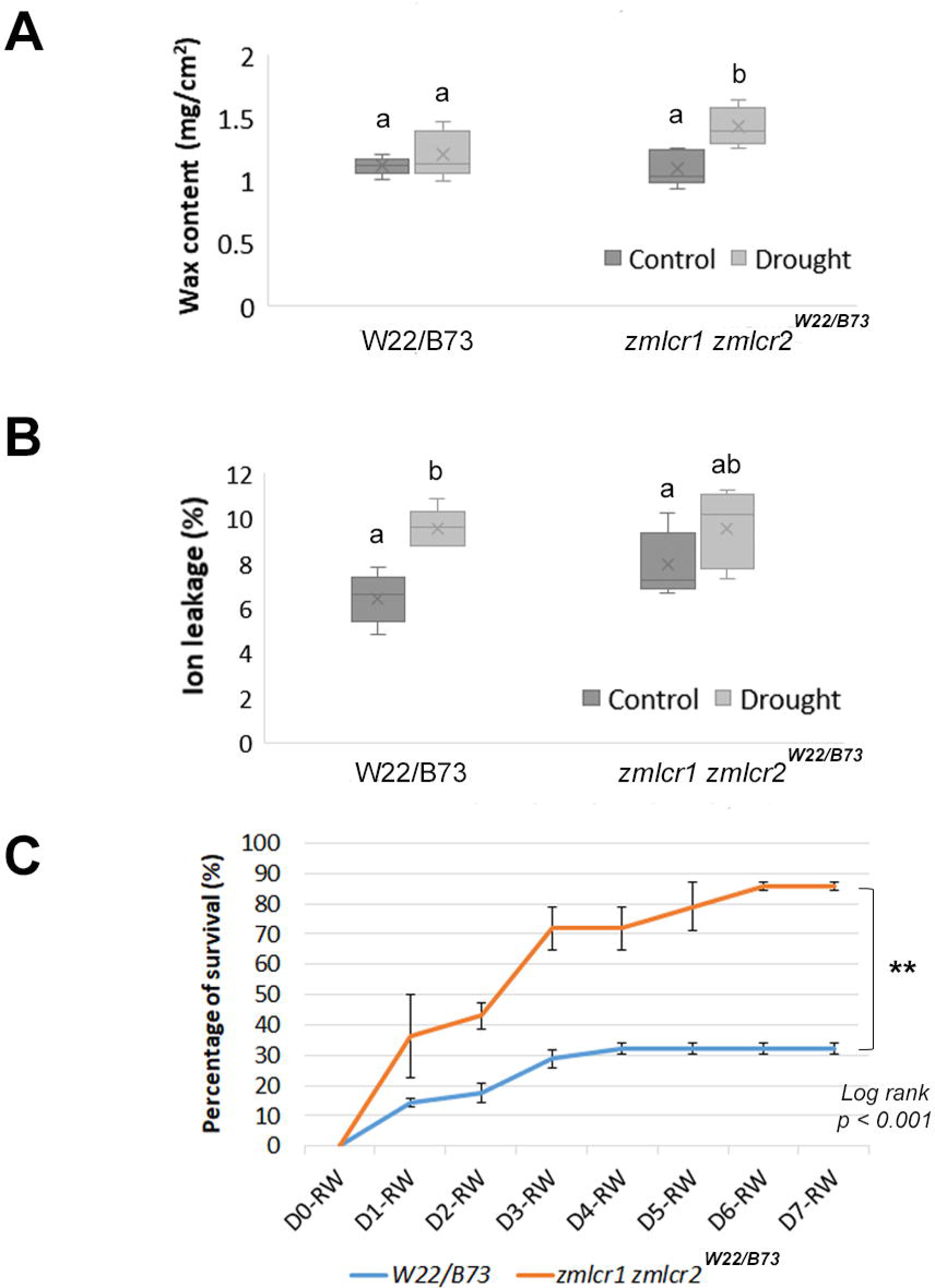
Wax content and membrane stability. A) Epicuticular wax content and B) Ion leakage of W22/B73 and *zmlcr1 zmlcr2^W22/B73^* hybrid seedlings under control and drought conditions. Different letters indicate significant differences according to ANOVA (p < 0.05), using genotype and treatment as factors, and Tukey’s post-hoc tests for multiple comparisons; n ≥ 5. C) Survival of W22/B73 and *zmlcr1 zmlcr2^W22/B73^* hybrid seedlings after 30 days of drought followed by re-watering during 7 days. Survival was assessed as the capability of plants to resume growth after re-watering. W22/B73 survival curve is different from *zmlcr1 zmlcr2^W22/B73^* hybrids, according to Kaplan–Meier log-rank analysis (n ≥ 13), followed by Holm-Sidak method for pairwise comparisons, ** = p < 0.001.

Next, in order to assess whether the beneficial effects in plant physiology, wax content and ion leakage observed in mutant hybrid seedlings result in a differential survival rate after a long period of drought, we allowed for a total of 30 days without irrigation and resume watering to determine plant survival. Our results indicate that *zmlcr1 zmlcr2^W22/B73^* mutant hybrids are more tolerant than W22/B73 hybrids, showing a higher capability of resuming growth during 7 days of re-watering (Fig. 2C). While only 32% of wild-type hybrids resumed growth after drought treatment, 85% of mutant hybrids survived the drought treatment (Fig. 2C), indicating that mutations in the hybrid background also results in higher survival rates, to a similar extent to that observed in the W22 inbred background (Miskevish et al., 2025).

### 3.3. Mutant hybrids have lower ROS production that wild-type hybrids

To investigate whether ROS production is affected in mutant hybrids and to evaluate oxidative stress in response to drought, we assessed H_2_O_2_ accumulation in leaves of wild-type and *zmlcr1 zmlcr2^W22/B73^* mutant hybrids using DAB staining (Fig. 3A). Under control conditions, W22/B73 leaves exhibited subtle DAB staining, indicating basal levels of H_2_O_2_, whereas staining was reduced in the mutant, as indicated by quantification of the DAB signal (Fig. 3A-B). Upon drought treatment, DAB staining intensified in wild-type leaves, consistent with elevated H_2_O_2_ levels (Fig. 3A); however quantitative analysis showed that the apparent increase in DAB staining observed in wild-type leaves under drought is not statistically different than staining in control conditions (Fig. 3B). Interestingly, hybrid mutants maintained low levels of staining (Fig. 3A), confirmed by the quantitative analysis showing lower DAB-stained area in the *zmlcr1 zmlcr2^W22/B73^*plants compared to wild type under both conditions (Fig. 3B). The reduced H_2_O_2_ accumulation in the mutants suggests a potential role of ZmLCR genes in regulating drought-induced oxidative stress.

**Figure 3:**
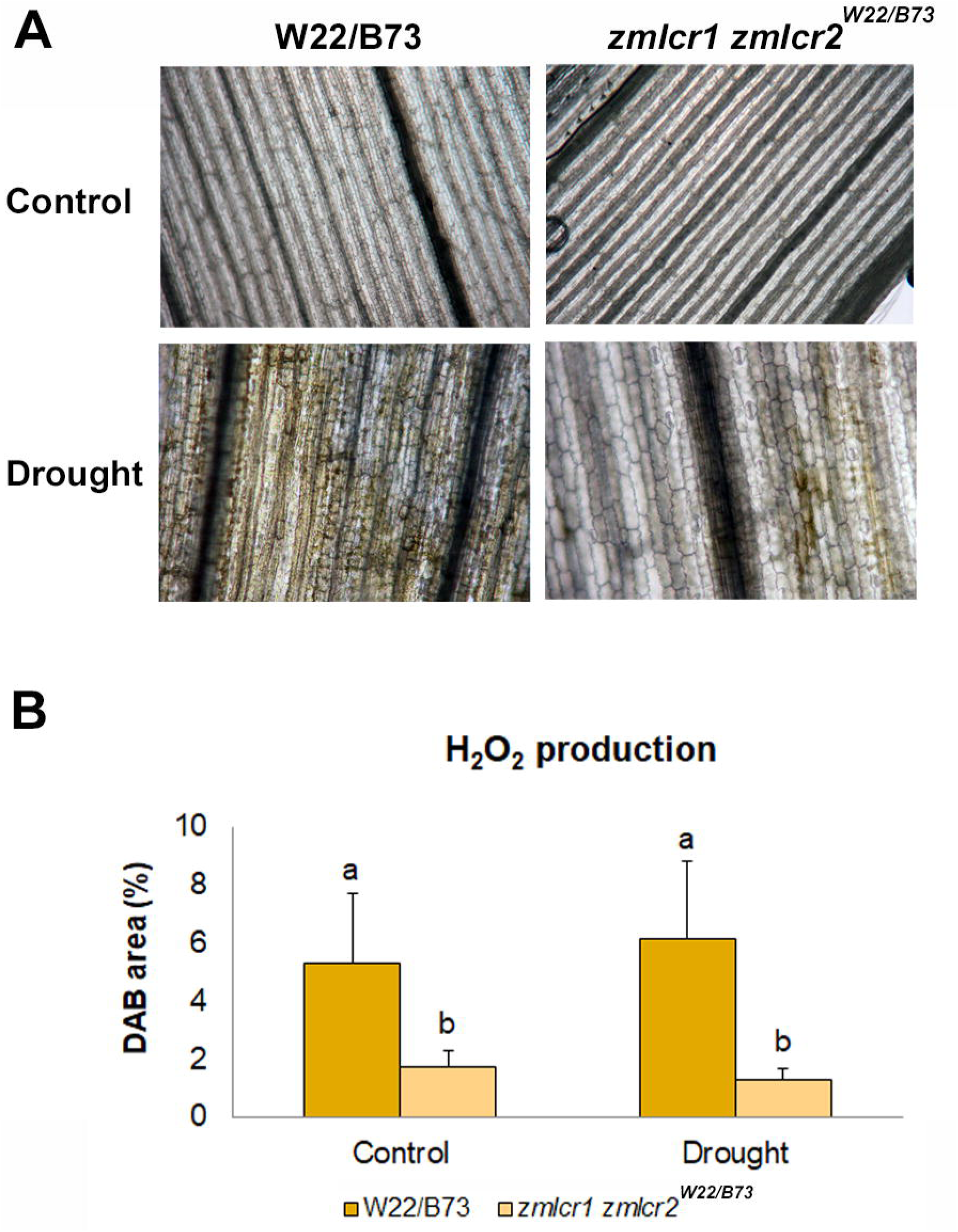
DAB staining of maize leaves to detect hydrogen peroxide accumulation. A) Representative images of DAB-stained leaves from wild-type (W22/B73) and *zmlcr1 zmlcr2^W22/B73^* mutant hybrids grown under normal irrigation (Control) or without irrigation for 15 days (Drought) conditions. DAB (3,3′-diaminobenzidine) staining indicates *in situ* accumulation of hydrogen peroxide (H_2_O_2_), visualized as brown precipitate. B) Quantification of DAB-stained area expressed as percentage of DAB area, calculated as total DAB pixels over total pixels in the corresponding channel. Five microscope photographs of DAB-stained leaves from each genotype and treatment were used for each analysis. The stained areas were identified by using the normalized threshold feature and processed using the vector H DAB from ImageJ Fiji. Mean ± SD is represented; different letters indicate significant differences according to ANOVA (p < 0.05; n ≥ 6), with genotype and treatment as factors, followed by Tukey’s post-hoc tests for multiple comparisons.

### 3.4. No physiological differences are observed in adult wild-type and mutant hybrids in field conditions

To characterize the physiological status of the plants grown in experimental field plots, we analyzed the same physiological parameters determined for seedlings, on 75-days-old wild-type and mutant hybrids. In this case, we also analyzed two different irrigation conditions, but instead of drought we evaluated rainfed, i.e. plants that only received rain water, since this is the watering regime typically used for this crop, which is characterized by scarce water availability in summer season and elevated temperatures, and we compared that condition to plants receiving supplemented irrigation, i.e. plants that were also manually watered every 2 days without rain, to mimic growing conditions in a more tropical setting with high water provision.

The statistical analysis of the measurements of the different parameters (E, gs, Pn, iWUE and greenness index) indicated that there is no interaction between the analyzed genotypes, W22/B73 and *zmlcr1/zmlcr2^W2/B73^*and the type of irrigation, rainfed or supplemented irrigation, for any of the parameters analyzed (Fig. 4). It was also determined that there are no differences in any of the parameters analyzed between wild and double mutant plants, under rainfed or supplemented irrigation conditions (Fig. 4), indicating that the physiology of mutant hybrids does not differ from that of wild-type plants, and mutations do not provide any detrimental or beneficial effects in the physiology of adult hybrids at this point in growth, when cultivated in field conditions.

**Figure 4:**
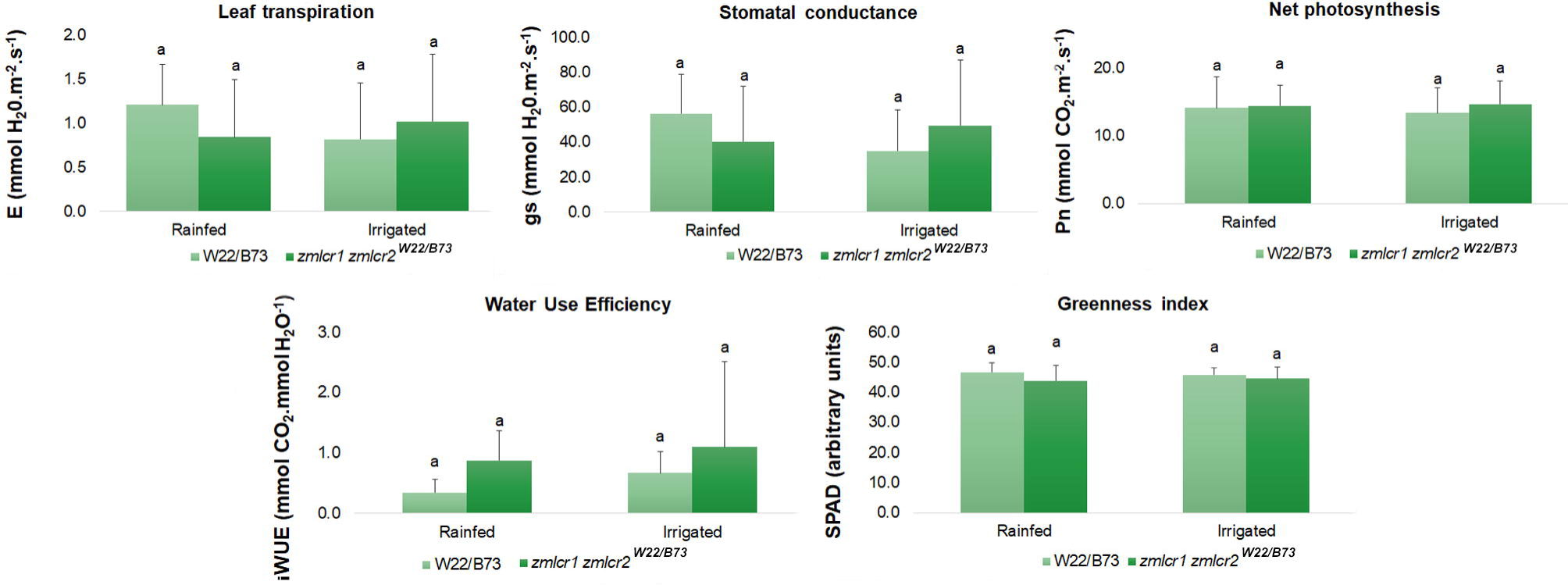
Characterization of W22/B73 and *zmlcr1 zmlcr2^W22/B73^* hybrid plants in experimental field plots. Physiological parameters determined in 75-days-old hybrid plants are shown: Greenness index, Leaf transpiration (E), stomatal conductance (gs), Net photosynthesis (Pn) and Water Use Efficiency (WUE) for plants grown under rainfed or irrigated conditions. Plant physiology is not affected in mutants under both watering regimes. Mean ± SD is represented; identical letters indicate there are no significant differences according to ANOVA (p < 0.05), using genotype and treatment as factors and Tukey’s post-hoc tests for multiple comparisons; n ≥ 10.

### 3.5. Nutritional composition is not affected in the mutants

In order to evaluate whether the mutations influence the nutritional content of the plants, we analyzed vegetative tissues from wild-type and mutant hybrids collected from the experimental plots after ear harvesting. Each evaluated parameter contributes to study different aspects of plant nutritional value, useful when formulating animal feed and to assess crop quality. Interestingly, analysis of dry matter (DM), crude protein (CP), acid detergent fiber (ADF), neutral detergent fiber (NDF), acid detergent lignin (ADL) and ether extract (EE) shows that mutant hybrids do not present differences in any of these parameters, indicating mutations are not beneficial or detrimental to the nutritional characteristics of the plants (Table 1).

**Table 1:** Nutritional content in W22/B73 and *zmlcr1 zmlcr2^W22/B73^* hybrids.

|  | W22/B73 | *zmlcr1 zmlcr2*<sup>W22/B73</sup> |
| --- | --- | --- |
| % DM | 83.56 ± 9.52<sup>a</sup> | 64.47 ± 14.64<sup>a</sup> |
| % CP | 2.48 ± 0.88<sup>a</sup> | 2.09 ± 0.47<sup>a</sup> |
| % ADF | 44.98 ± 6.40<sup>a</sup> | 45.56 ± 2.22<sup>a</sup> |
| % NDF | 66.49 ± 7.22<sup>a</sup> | 68.72 ± 4.62<sup>a</sup> |
| % ADL | 5.47 ± 0.79<sup>a</sup> | 5.51 ± 0.38<sup>a</sup> |
| % EE | 0.96 ± 0.20<sup>a</sup> | 0.679 ± 0.13<sup>a</sup> |

<sup>a</sup>: no statistical difference between genotypes (Student's T-test, p < 0.05, n = 3)

### 3.6. Wild-type and mutant hybrids show no difference in flowering time

To continue with the characterization of hybrids in the experimental plot, the flowering time of W22/B73 and *zmlcr1 zmlcr2^W22/B73^*hybrids was determined as the number of days until the appearance of stigmas in the female inflorescence (stage R1) and the time of anther appearance and pollen production was also analyzed, under the two irrigation conditions mentioned above. The statistical analysis established that there are no differences in flowering time between wild-type and double mutants hybrids under rainfed or supplemented irrigation conditions (Fig. 5A). In addition, wild-type hybrids flower at the same time under both watering conditions, but *zmlcr1 zmlcr2^W22/B73^* mutant plants respond significantly to the supplemented watering condition, flowering slightly earlier (Fig. 5A). Under rainfed conditions, *zmlcr1 zmlcr2^W22/B73^* hybrids flowered on average at 72.9 days, while under supplemented irrigation the average flowering time was 71.2 days (Fig. 5A). Regarding anther appearance and pollen production, we observed no statistically significant differences, with an average of 72.2 and 73.0 days for W22/B73 and *zmlcr1 zmlcr2^W22/B73^*, respectively in rainfed conditions, in comparison to 71.8 and 71.5 for wild-type and mutant hybrids in supplemented irrigation (Fig. 5C).

**Figure 5:**
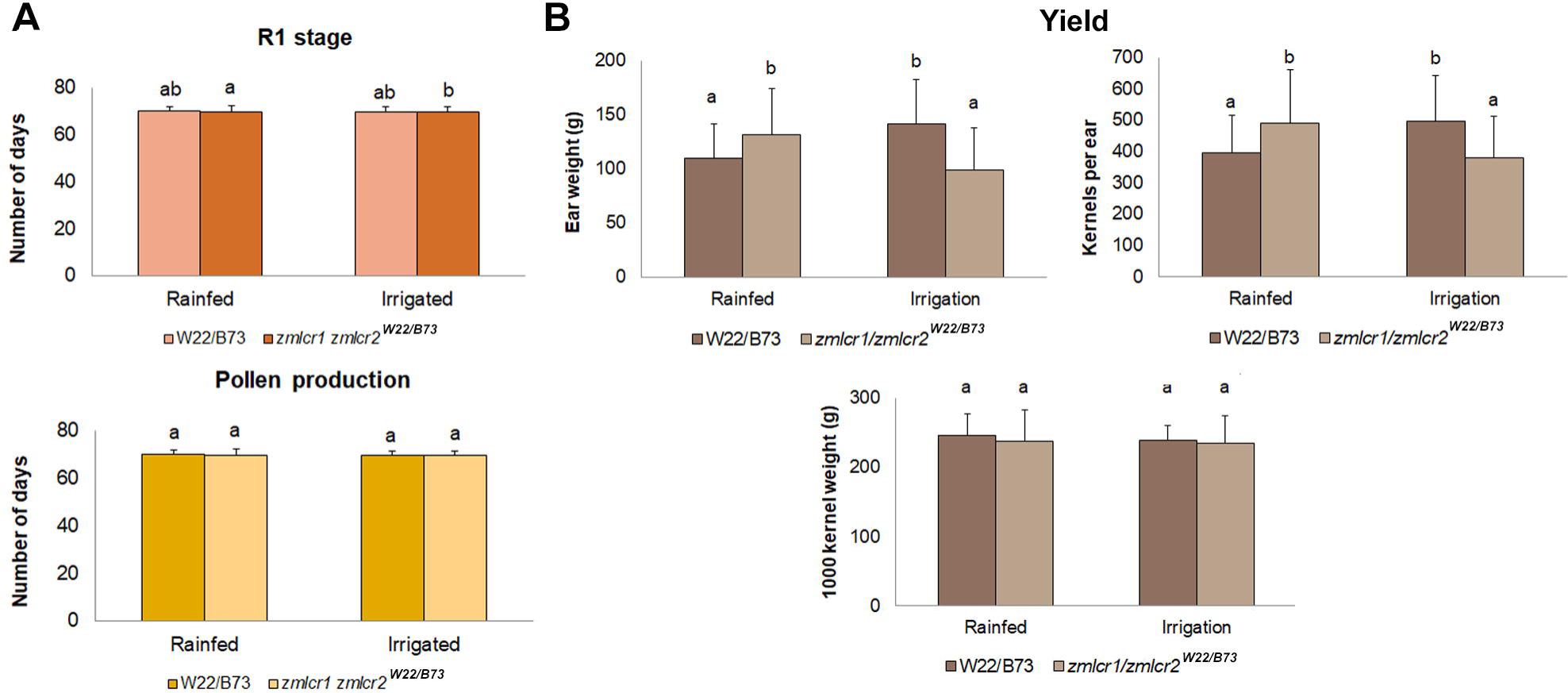
Flowering and yield in W22/B73 and *zmlcr1 zmlcr2^W22/B73^* hybrid plants in experimental field plots. A) Days until the appearance of stigmas (stage R1) and appearance of anthers and pollen production in hybrid plants under rainfed and supplemented irrigation conditions. Mean ± SD is represented; different letters indicate significant differences according to two-factor ANOVA (p < 0.05), followed by multiple comparisons by Tukey’s method; n ≥ 30. B) Ear weight, kernels per ear and 1000 kernel weight of wild-type and mutant hybrids, under rainfed conditions and supplemented irrigation. Mean ± SD; different letters indicate significant differences according to two-factor ANOVA (p < 0.05), followed by multiple comparisons by Tukey’s method; n ≥ 10.

### 3.7. Yield is increased in hybrid mutants under rainfed conditions

To analyze whether the mutations in *ZmLCR* genes have an influence on the yield of hybrid plants, we analyzed yield parameters of plants grown in rainfed conditions, similar to those of typical agricultural production in our country location, and on plants that received supplemental irrigation. Once the plants grown on experimental plots reached stage R6, after 125 days of growth, ears were harvested manually and dried at 27 °C for one week.

We determined that ear weight and the number of kernels per ear were higher for *zmlcr1 zmlcr2^W22/B73^* in rainfed conditions, but lower for mutant hybrids that received supplemented irrigation (Fig. 5B). Ears from wild-type rainfed hybrids weighed 110.00 g and had 396.28 kernels on average, whereas ears from mutant hybrids weighed 131.79 g and had 490.85 kernels (Fig. 5B). Conversely, the weight of 1000 kernels did not show statistical differences between both genotypes and watering conditions, with average values close to 240 g in all cases, indicating that kernel weight is not affected (Fig. 5B).

## 4. Discussion

Traditional plant breeding and hybrid development have long served as essential strategies to improve agronomic traits such as yield, pest resistance, and stress tolerance. More recently, genetic engineering has enabled the precise introduction or alteration of genes, generating genetically modified crops with novel characteristics (Caradus, 2023). However, regardless of the approach, integrating desirable traits often carries unintended consequences, commonly referred to as penalties (Cellini et al., 2004). Since commercial maize production predominantly relies on hybrid lines, evaluating the impact of *zmlcr1/zmlcr2* mutations within a hybrid context is particularly relevant to real-world agricultural scenarios. Our study demonstrates that introducing these mutations into a hybrid genetic background enhances seedling physiological performance under normal watering conditions and in drought stress, as well as increasing yield in rainfed field conditions, without evident physiological penalties, but reducing yield in an environment with excess water availability.

Abiotic stress is of utmost importance in agronomy, exerting significant negative effects on plant development by disrupting morphological, physiological, biochemical, and molecular processes. These stresses often impair critical stages such as flowering and reproductive transitions, directly affecting crop yield. In light of climate change and the increasing frequency of extreme weather events, the urgency to develop resilient crops has intensified (Basso et al., 2019; Leakey et al., 2019; Kim et al., 2019; Chaudhry and Sidhu, 2022).

In our analysis of *zmlcr1 zmlcr2^W22/B73^* double mutant hybrid seedlings, we observed enhanced physiological performance even under well-watered conditions, contrasting with previous findings in inbred W22 mutants, where no such differences were noted under similar conditions (Miskevish et al., 2025). In hybrids, mutant seedlings showed improved leaf-level traits including reduced transpiration, increased stomatal conductance and higher net photosynthesis, ultimately leading to improved intrinsic water use efficiency (iWUE). These results suggest that the hybrid genetic context amplifies the physiological advantages conferred by ZmLCR mutations, even under normal watering conditions.

Plants deploy diverse biochemical and physiological adaptations to cope with the lack of water, involving complex regulatory networks (Tekle & Alemu, 2016). Under drought conditions, our hybrid mutants exhibited superior tolerance compared to wild-type counterparts, as evidenced by higher iWUE and net photosynthesis, increased epicuticular wax content, and reduced membrane damage. While these findings align with prior results in inbred lines, the notable distinction is that hybrid mutants also demonstrated benefits under non-stressed conditions (Miskevish et al., 2025). Additionally, mutant hybrids showed a high recovery rate (∼85%) after prolonged drought, similar to inbred mutants, further reinforcing the value of these genetic modifications. The observed increase in epicuticular wax likely contributes to reduced water loss, supporting both drought resilience and water use efficiency. Interestingly, membrane stability under drought, as measured by ion leakage assays, improved significantly in the hybrid background, an effect not seen in inbred lines, highlighting an enhanced drought-response phenotype in hybrids. This result could also be connected to the observed H_2_O_2_ production in mutant hybrids, where lower levels of reactive oxygen species could be beneficial to maintain membrane integrity even under stress conditions. Moreover, it has been shown that when maintained at relatively low levels, ROS act as signaling molecules that regulate plant growth, development, and adaptation to adverse conditions (Wang et al., 2024), which could also be a factor supporting the increased survival for mutant hybrids after drought treatment.

The most drought-sensitive stages in maize development are the vegetative phase near flowering and the grain-filling period (Sah et al., 2020; Sheoran et al., 2022), while this crop is especially sensitive to excess moisture in early vegetative and tasseling stages (Zaidi et al., 2004). To evaluate the impact of ZmLCR mutations under production-like conditions, we assessed mutant and wild-type hybrids in field plots under rainfed conditions, with some plots receiving supplemental irrigation. The goal was to mimic real agricultural settings which are typically characterized by high temperatures and limited rain water availability during summer season in our field location in Argentina, and assess potential yield implications in comparison to a simulated setting of increased rain, accomplished by the manually added irrigation. Notably, no major physiological differences were observed between mutant and wild-type hybrids in these plots, potentially due to the timing of the measurements, which were taken during reproductive stages when stress effects are less pronounced, and to more moderate water limitation in field conditions compared to controlled drought treatments used in growth chambers (Miskevish et al., 2025). These results suggest that the observed increase in iWUE represents an advantage in mutants especially under severe drought conditions and particularly during early plant development, but is not evidenced in adult plants after flowering.

No significant differences in flowering time were found between wild-type and mutant hybrids, consistent with prior studies indicating that miR394 regulates flowering in *Arabidopsis thaliana* independently of LCR, the only known miR394 target in that species (Bernardi et al., 2022). Future studies targeting *zma-MIR394* genes would help determine whether the regulatory role of miR394 is conserved in maize. For now, our results support the conclusion that ZmLCR genes do not influence flowering time in maize. In addition, and taking into consideration that our study concludes that mutations do not affect the nutritional content of hybrid plants, taken together all these findings suggest that the mutations do not adversely affect key physiological or developmental processes in hybrid plants, supporting their agronomic potential.

A particularly compelling finding is the observed increase in yield among mutant hybrids, indicated by greater ear weight and more kernels per ear compared to W22/B73 controls. This enhancement may be a consequence of the improved seedling physiology and water use efficiency during vegetative growth, which positively influences reproductive success. Although lower yields were registered for plants grown in plots with supplemented irrigation, suggesting limited suitability in regions with excess moisture, these results highlight the strong potential of these mutants for improving maize productivity in rainfed and in water-limited environments.

## 5. Conclusion

Our findings position ZmLCR genes as promising targets for maize genetic improvement, especially in hybrid backgrounds. Mutant lines exhibit improved physiological parameters and reduced ROS production at the seedling stage, under both normal or drought conditions. Additionally, under drought stress, these mutants show increased epicuticular wax accumulation, contributing to enhanced survival after prolonged water deprivation. Importantly, hybrid mutants cultivated in the field, under production-like conditions maintain normal flowering time and nutritional composition, while achieving a notable increase in yield. These findings pave the way for developing higher-yielding, drought-tolerant maize cultivars without compromising essential agronomic traits.

### Author contributions

MD and GD were responsible for the study conception and design. Material preparation, data collection and analysis were performed by FB, PGP, CJ and SW. ID contributed to experimental design and management of plant growth and care in experimental plots and JM performed the nutritional analysis. The first draft of the manuscript was written by MD and all authors commented on initial versions of the manuscript. All authors read and approved the final manuscript.

## Data Availability

Data generated and/or analyzed during the current study are included in the manuscript or as supplemental materials.

## Acknowledgements

Authors would like to thank Marcos Reyes and Mauro Alisio, CPA members of the Argentinean Consejo Nacional de Investigaciones Científicas y Técnicas (CONICET) for collaboration with plant care and maintenance. MD, GD and ID are members of the Researcher Career of CONICET and FB is a CONICET doctoral fellow.

